# Antagonistic interactions between odorants alter human odor perception

**DOI:** 10.1101/2022.08.02.502184

**Authors:** Yosuke Fukutani, Masashi Abe, Haruka Saito, Ryo Eguchi, Toshiaki Tazawa, Claire A. de March, Masafumi Yohda, Hiroaki Matsunami

## Abstract

The olfactory system detects a vast number of odorants using hundreds of olfactory receptors (ORs), the largest group of the G protein-coupled receptor (GPCR) superfamily. Each OR is activated by specific odorous ligands. Like other GPCRs, activation of ORs may be blocked through antagonism. Recent reports highlight widespread antagonisms in odor mixtures influencing olfactory neuron activities. However, it is unclear if and how these antagonisms influence perception of odor mixtures. Here we show that odorant antagonisms at the receptor level alter odor perception. Using a large-scale heterologous expression, we first identified a set of human ORs that are activated by methanethiol and hydrogen sulfide, two extremely potent volatile sulfur malodors. We then screened odorants that block activation of these ORs and identified a set of antagonists, including β-ionone. Finally, human sensory evaluation revealed that odor intensity and unpleasantness of methanethiol were decreased by β-ionone. Odor intensity of β-ionone itself is not correlated with the degree of suppression of malodor sensation. Suppression was also not observed when methanethiol and β-ionone were simultaneously introduced to different nostrils. Together, our data supports the model that odor sensation is altered through antagonistic interactions at the level of the ORs.

## Introduction

Animals detect and discriminate numerous environmental odorants through combinatorial activation of membrane receptors expressed in olfactory sensory neurons (OSNs). In humans, the predominant receptors are the olfactory receptors (ORs), which are the largest protein-coding gene family of Class A G-protein coupled receptors (GPCRs) with ∼400 members ^1-5^.

When an odorous agonist binds to an OR, the OR favors an active conformation that initiates a signal transduction cascade, leading to depolarization of the OSNs. Each OSN expresses only one type of OR, meaning that the activation of an OSN is determined by the activation of the OR ^6,7^. A single OR can respond to various odorants, and one odorant activates multiple ORs. Thus, the combinatorial pattern of OR activation is foundational for the detection and discrimination of a given odor ^8-11^.

In addition to acting as an agonist, comparative to ligands for canonical GPCRs, some odorants can function as antagonists or inverse agonists. By binding to ORs, odorants can inhibit the activation by agonists in a competitive manner or stabilize the receptor in the inactive conformation. Previous *in vivo* and *ex vivo* studies as well as *in vitro* research using heterologously expressed ORs have shown that odorants are often capable of acting as antagonists for ORs ^12-19^. Natural odors are usually mixtures of odorants, so antagonistic interactions between odorants at ORs are likely. Mixture suppression, in which odor intensity is less than a linear sum of component odorants, is commonly observed ^20,21^. Previous studies highlighted the role of central olfactory processing in mixture suppression^22-24^. Yet peripheral events, such as antagonistic interactions of odorants at the ORs, were also proposed as an underlying mechanism of mixture suppression ^24,25^. Despite widespread antagonism on OSNs in odor mixtures, there is scare evidence supporting a role of odorant antagonism in odor perception ^20,26^. This is partly because OR antagonists also act as agonists for other ORs, making it challenging to distinguish the role of antagonism in odor perception from central processing.

In this study, we addressed the effects of odorant antagonism in odor perception by focusing on a set of ORs that are activated by extremely potent and unpleasant volatile sulfur compounds (VSCs), including methanethiol (methyl mercaptan, CH_3_SH). Antagonists, such as β-ionone, blocked the VSC OR response and modified the sulfur odorants odor perception.

## Results

### Screening human ORs that respond to gaseous sulfur compounds

Previously, we developed a method to detect *in vitro* OR responses to odors in the vapor phase ^11,27^. Using this technique with modifications (see Methods), we performed an OR screening assay for two highly volatile sulfur compounds (VSCs), methanethiol and hydrogen sulfide (Fig. 1A and Extended data Fig. 1).

**Figure 1.**
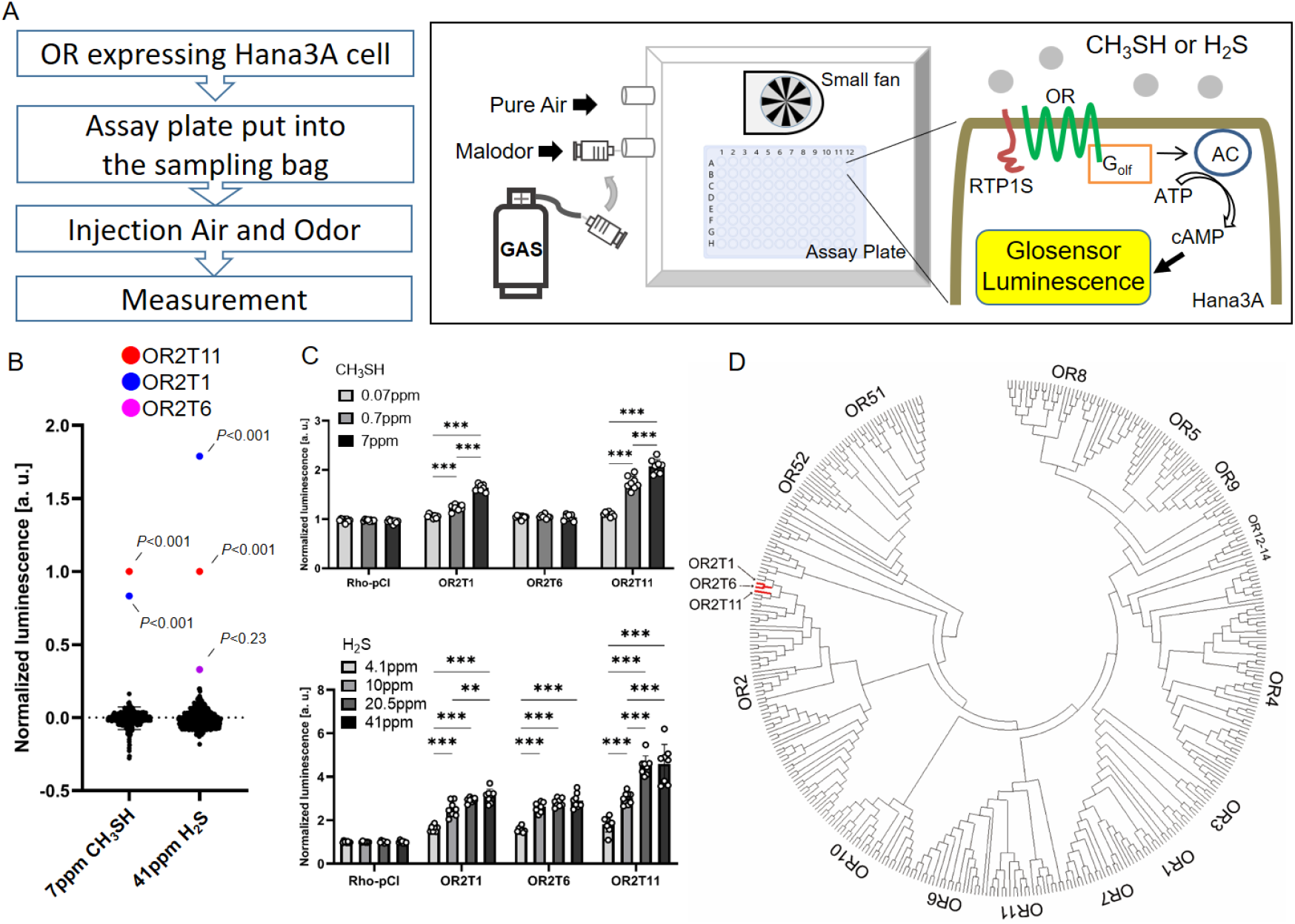
Screening of human ORs responding volatile sulfur compounds. A) Schematic representation of the vapor stimulation assay with the OR signal transduction pathway. AC; adenylyl cyclase, ATP; adenosine triphosphate, cAMP; cyclic adenosine monophosphate, RTP1S; receptor transporting protein 1 short. B) Response of 359 unique human ORs against 7 ppm CH_3_SH and 41 ppm H_2_S in vapor phase. Multiple comparisons were performed using one-way analysis of variance (ANOVA) followed by Dunnett’s Test. C) Vapor dose response analysis of identified human ORs. The normalized luminescence value indicates the S/N ratio before and after stimulation of CH_3_SH or H_2_S. Multiple comparisons were performed using one-way analysis of variance (ANOVA) followed by Dunnett’s Test (*p<0.05. **p<0.01, ***p<0.001). D) Similarity of OR2T1, OR2T6 and OR2T11 in phylogenic tree of human ORs.

By screening the response of 359 human ORs, representing the vast majority of intact ORs encoded on the human genome, to methanethiol and hydrogen sulfide, we identified several candidate hits that exhibited strong responses (*p*<0.001, one-way ANOVA followed by Dunnett’s test) (Fig. 1B and Supplementary Data). A secondary screening confirmed concentration-dependent responses of OR2T11 and OR2T1 to methanethiol and hydrogen sulfide as well as the response of OR2T6 to hydrogen sulfide (Fig. 1C). These three ORs share relatively high amino acid sequence similarities among the OR family members (Identities; 73% (222/305), 66% (206/310), and 63% (193/304) in OR2T1/OR2T6, OR2T1/OR2T11, OR2T6/OR2T11, respectively) (Fig. 1D).

OR2T11 was previously shown to respond to various thiol molecules in a copper- and silver ion-dependent manner ^28-30^. We examined the metal dependence of the ORs’ responses to methanethiol and hydrogen sulfide. Consistent with the previous reports, the addition of copper in the media dramatically enhanced the response to both VSCs. The addition of silver produced moderate response enhancement, while the addition of zinc showed no effect (Extended data Fig. 2A). The effect of copper was diminished by the addition of the copper chelator tetraethylenepentamine (TEPA) (Extended data Fig. 2B). Our data supports the previously suggested model that metal-sulfur complexes activate sulfur-responsive ORs.

### Identification of antagonists

Antagonists or inverse agonists of ORs that are potently activated by the sulfur odorants may block the perception of their unpleasant odor, potentially leading to the development of novel deodorants. To identify antagonists that block the activity of VSC-responding ORs, we screened the effects of 100 odorants on OR2T11 response to methanethiol and hydrogen sulfide. OR2T11-expressing cells in a buffer containing each compound at a final concentration of 100 µM were stimulated by 7 ppm methanethiol and 41 ppm hydrogen sulfide. A subset of ketones, including specific ionones and damascones, showed inhibitory effects on the OR responses against the tested VSCs (Fig. 2A). β-ionone was the most potent antagonist among the tested odorants (LogIC50 = -4.85) (Fig. 2B and Extended data Fig. 3A) and did not alter the activity of OR2T11 by itself when tested without VSCs (Fig. 2C). In contrast, Iso E super (tetramethyl acetyloctahydronaphthalen)^31,32^, which is one of the most commonly used ketone fragrances for deodorant, did not show potent antagonistic effects on OR2T11 (Fig. 2B).

**Figure 2.**
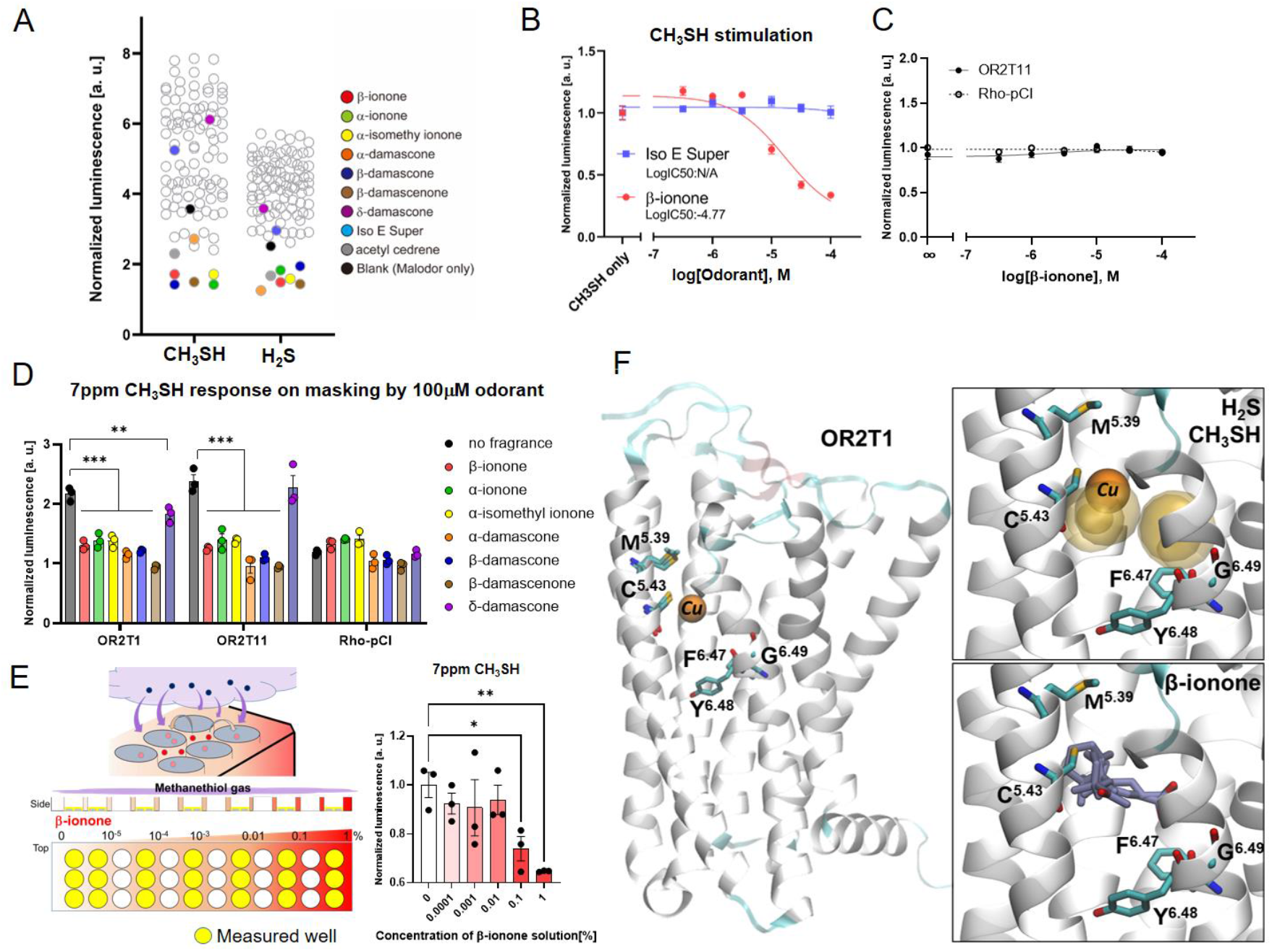
Antagonists of VSC responding human ORs. A) Screening for 100 odorants that inhibit the VSC response of OR2T11. B) Dose response inhibition of β-ionone (red) and Iso E Super (blue) against OR2T11 with the vapor concentration of CH_3_SH held constant at 7ppm. IC50 value was calculated using Graph pad Prism software. C) Dose−response curves of OR2T11 against β-ionone. D) Antagonistic effect of damascone & ionone analogs. 7ppm CH_3_SH stimulation of human OR2T1 and OR2T11 masked by 100µM odorants. Multiple comparisons were performed using one-way analysis of variance (ANOVA) followed by Dunnett’s multiple comparison test (\*\**p*<0.01, \*\*\**p*<0.001). E) Inhibitory effect in vapor phase. Soon after adding 25 µL of β-ionone (red) solution between the wells of the 96-well plate, OR2T11 expressing cells were quickly stimulated by CH_3_SH gas (blue). The normalized luminescence value indicates the S/N ratio before and after stimulation of CH_3_SH. Multiple comparisons were performed using one-way analysis of variance (ANOVA) followed by Dunnett’s Test (\**p*<0.05. \*\**p*<0.01) F). Left. View of the OR2T1, 2T6 and 2T11 Alphafold models. Residues M^BW5.39^ and C^BW5.43^ are represented for the three ORs while the rest of the structure is only shown for OR2T1 for clarity. The docked copper is shown in orange Van der Waals volume. The toggle switch of mammals ORs, the FYG motif at positions F^BW6.47^ to G^BW6.49^, is also developed and serves as reference for the bottom of the odorant binding site. Right. Top. All the docking position of H_2_S and methanethiol in the binding cavity of the three OR2T are represented in transparent volume. Bottom. The docking position of β-ionone in the three OR2T binding cavity are represented in blue licorice.

Next, we examined the inhibitory effects of the ionone and damascone molecules on the responses of OR2T11, OR2T1, and OR2T6 to the VSCs. We observed that all tested ORs were similarly inhibited by a given antagonist, suggesting a common blocking mechanism. (Fig. 2D and Extended data Fig. 3B). The possibility that the inhibition of the odorants was caused by adverse effects on the assay system, such as cytotoxicity, was excluded because another OR, mouse Or2aj6 (also known as Olfr171 and MOR273-1), was activated by these compounds in the same assay (Extended data Fig. 3C). In the vapor stimulation assays, a mixture of methanethiol and β-ionone also showed a concentration-dependent and molecule-specific inhibitory effect against OR2T11 and OR2T6 (Fig. 2E and Extended data Fig. 4A). Since the concentration of methanethiol did not decrease by mixing with β-ionone in the sampling bag (Extended data Fig. 4B), chemical reactions between methanethiol and β-ionone is unlikely to be a cause of suppression of OR activations.

To gain insight into mechanism underlying the antagonisms, we performed docking of the ligands on the structural models of OR2T families based on AlphaFold2 ^33^ (Extended data Fig. 5). Following previous publications and our data suggesting copper as an essential co-factor to antagonism (Extended data Fig. 2), OR bound to sulfur odorants by coordinating with a copper cation at the level of sulfur residues ^28-30,34,35^. The docking of copper resulted in a binding close to C^BW5.43^ and M^BW5.39^. These residues, conserved in a small subset of ORs including OR2T1, OR2T6, and OR2T11, pointed towards the odorant binding cavity (Fig. 3F and Extended data Fig. 6A). The bottom of this cavity was defined by the OR toggle switch, the “FYG” motif (BW6.47 to BW6.49), in the transmembrane helix (TM) 6 ^36^. Hydrogen sulfide and methanethiol were docked on the copper bound OR structures. All poses were located in between the copper and the toggle switch (Fig 3F and Extended data Fig. 6), suggesting a possibility of effective binding for these two molecules in OR2T1, OR2T6, and OR2T11. The β-ionone binding location overlapped with that of the Cu-sulfur odorant in all three structures when it was docked into the cavity of OR2T1, OR2T6, and OR2T11. This model is consistent with the idea that β-ionone antagonizes OR activation by competitively blocking the binding of Cu-sulfur odorant.

**Figure 3.**
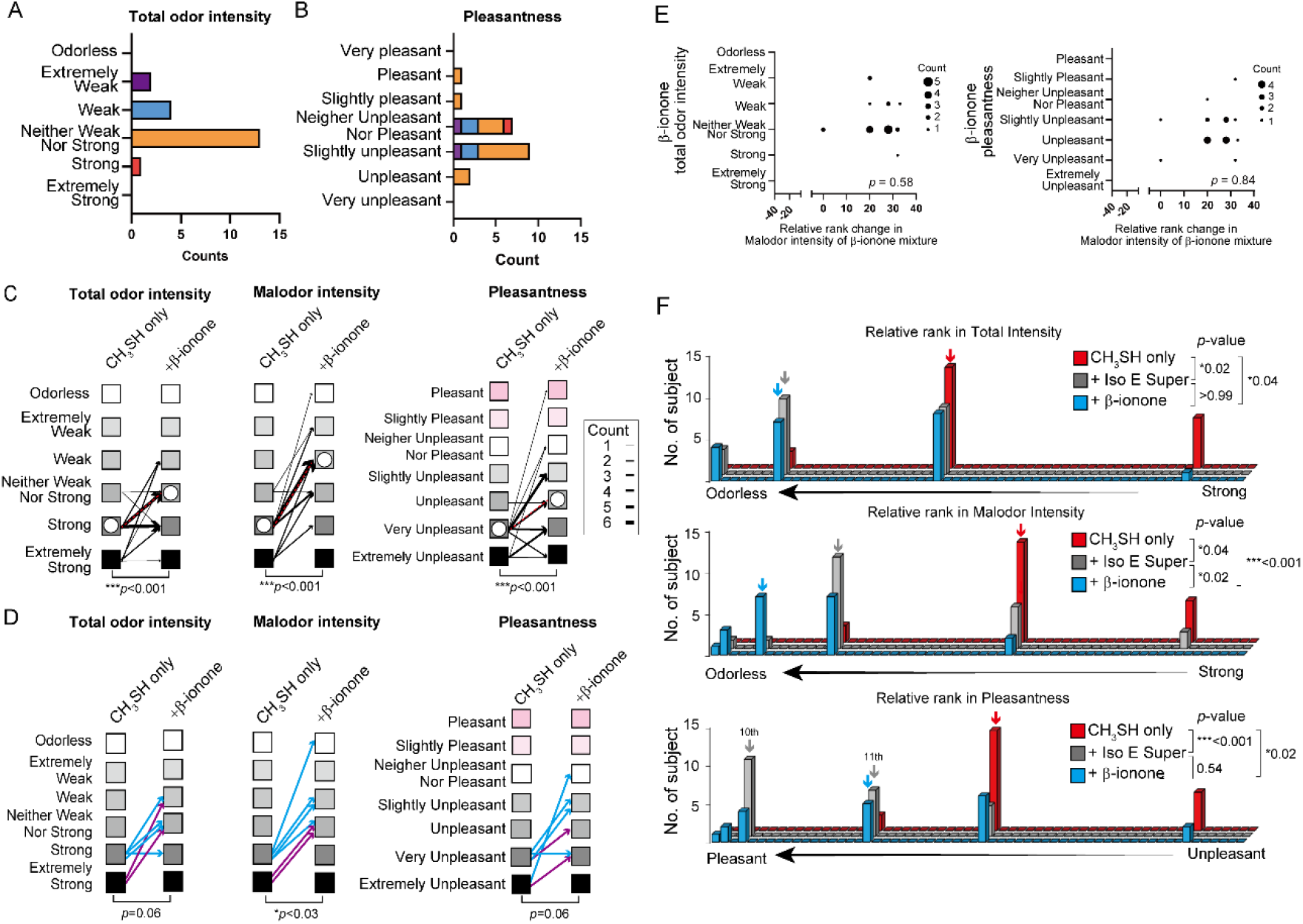
β-ionone changes odor perception against VSC. Human Sensory evaluation test against CH_3_SH gas containing β-ionone, an effective antagonist. A) Odor intensity value of β-ionone evaluated by 20 subjects. B) Pleasantness value of β-ionone. Color matched the odor intensity score of each subjects representing in Fig 3A. Comparison test was performed using the non-parametric Wilcoxon test Test (\*\*\**p*<0.001). C) Results of evaluation test Total odor intensity Malodor intensity and Pleasantness. Line thickness: The number of subjects who made the same evaluation transition in mixture gas. Median showed the white circle in each condition. D) Results of evaluation test limited to subjects who is difficult to feel β-ionone, Blue :Weak, purple:Extremely weak in Fig. 3A. Each line indicates one subject. Comparison test was performed using the non-parametric Wilcoxon test Test (\**p*<0.05). E) Correlation analysis between relative rank change in malodor intensity by mixing with β-ionone (X-axis) and total odor intensity (Upper) or pleasantness (Lower) score of the β-ionone (Y-axis). Plot size: number. Correlation values were calculated by nonparametric Spearman correction test. F) Relative rank in Total intensity, malodor intensity and pleasantness. Median showed the color arrows in each condition. Both are shown when the 10th and 11th are different. Multiple comparison test was conducted using nonparametric Friedman’s test (\**p*<0.05)

### Human sensory evaluation test

Thus far, we have identified that OR2T family members robustly respond to the tested VSCs and that certain ketones, including β-ionone, act as antagonists. If antagonists inhibit the response of all receptors that respond to particular VSCs, it is expected that inhibition of olfactory perception will occur. To test whether the antagonistic interactions affect olfactory perception, we designed a set of human sensory evaluation tests aiming to distinguish antagonistic effects and central processing in mixture suppression. We selected β-ionone (floral/violet/woody smell), the most potent antagonist we identified, in parallel with Iso E Super (woody/floral/amber/violet smell), which does not show potent antagonistic effects (Fig. 2B). While perceived odor intensities of β-ionone and Iso E Super varied among subjects, we adjusted their concentrations so that the intensities were matched as a group (Fig. 3A and Extended data Fig. 7 and 8). Iso E Super is more pleasant than β-ionone at the adjusted concentrations (\*\*\**p*<0.001, non-parametric Wilcoxon test) (Extended data Fig. 8B). Sensory evaluation tests were conducted by asking subjects to rate total odor intensity, malodor intensity, and pleasantness of methanethiol and its mixtures (methanethiol and β-ionone, methanethiol and Iso E Super). The subjects were separated into two groups with different evaluation orders (Extended data Fig. 8C). In order to clarify the evaluation criteria for the subjects, we presented the odors in a non-random order. We first conducted a non-blind test for methanethiol and asked subjects to score one value for both total odor intensity and malodor intensity. Subsequently, two odor mixtures (methanethiol with either β-ionone and methanethiol with Iso E Super) were presented in a random order to separately evaluate total odor intensity, malodor intensity, and pleasantness in the blind condition (Extended data Fig. 8C). Regardless of the order of odor presentation, the subjects’ evaluations were similar, excluding the role of odor adaptation in our tests (*p*>0.05, Mann-Whitney tests) (Extended data Fig. 7B). The total odor intensity of methanethiol and β-ionone mixtures was less than those of the methanethiol only, indicating mixture suppression (*p*<0.001, Wilcoxon test) (Fig. 3C Left). Additionally, the malodor intensity and unpleasantness of methanethiol was significantly reduced by mixing it with β-ionone (*p*<0.001, Kruskal-Wallis test) (Fig. 3C Center and Right). To test whether odor intensity of β-ionone alone predicts its suppressive effects on methanethiol, we first focused on the group with low odor intensity against β-ionone (6 out of 20 subjects) and found that the suppressing effect was evident (*p*=0.03, nonparametric Wilcoxon multiple comparison test) (Fig. 3D). When we compare the suppressing effects between this group and the group with higher odor intensity against β-ionone (14 out of 20 subjects), there was no statistically significant difference in methanethiol suppression (*p*=0.98, Mann-Whitney test). Moreover, both of the odor intensities and the pleasantness of β-ionone showed no significant correlation with the degree of suppressing effects against methanethiol (*p*=0.58 and 0.84, respectively, Spearman’s tests) (Fig. 3E). Together, the data suggests that odor intensity of β-ionone alone is independent from it’s odor suppressing effects. In the Relative rank, mixture of β-ionone was significantly higher than one of Iso E Super in the evaluation of malodor intensity (*p*=0.02, nonparametric Friedman’s multiple comparison test) (Fig. 3F). Iso E Super also reduced total odor intensity, malodor intensity, yet it was less effective in suppressing malodor intensity than β-ionone (Fig. 3F and Extended data Fig. 8).

Similarly, damascones (α-damascone, β-damascone, δ-damascone) and β-damascenone that inhibited OR response to the VSCs were effective in reducing the odor intensity and unpleasantness to methanethiol (Extended data Fig. 9). δ-damascone, which showed a low inhibitory effect in *in vitro* assay (Fig. 2D), also had the lowest suppressing effect among the tested antagonists in the sensory evaluation test. Together, the data is consistent with our hypothesis that antagonistic effects on ORs change odor perception.

### Nostril-specific stimulation

Finally, to differentiate the impact of peripheral antagonism vs central processing in the malodor suppression, we compared the impact of β-ionone on the perception of methanethiol when β-ionone was inhaled via the same versus a different nostril (Fig. 4A). The subjects were unable to distinguish which nostril the tested odor was presented to (Fig. 4B).

**Figure 4.**
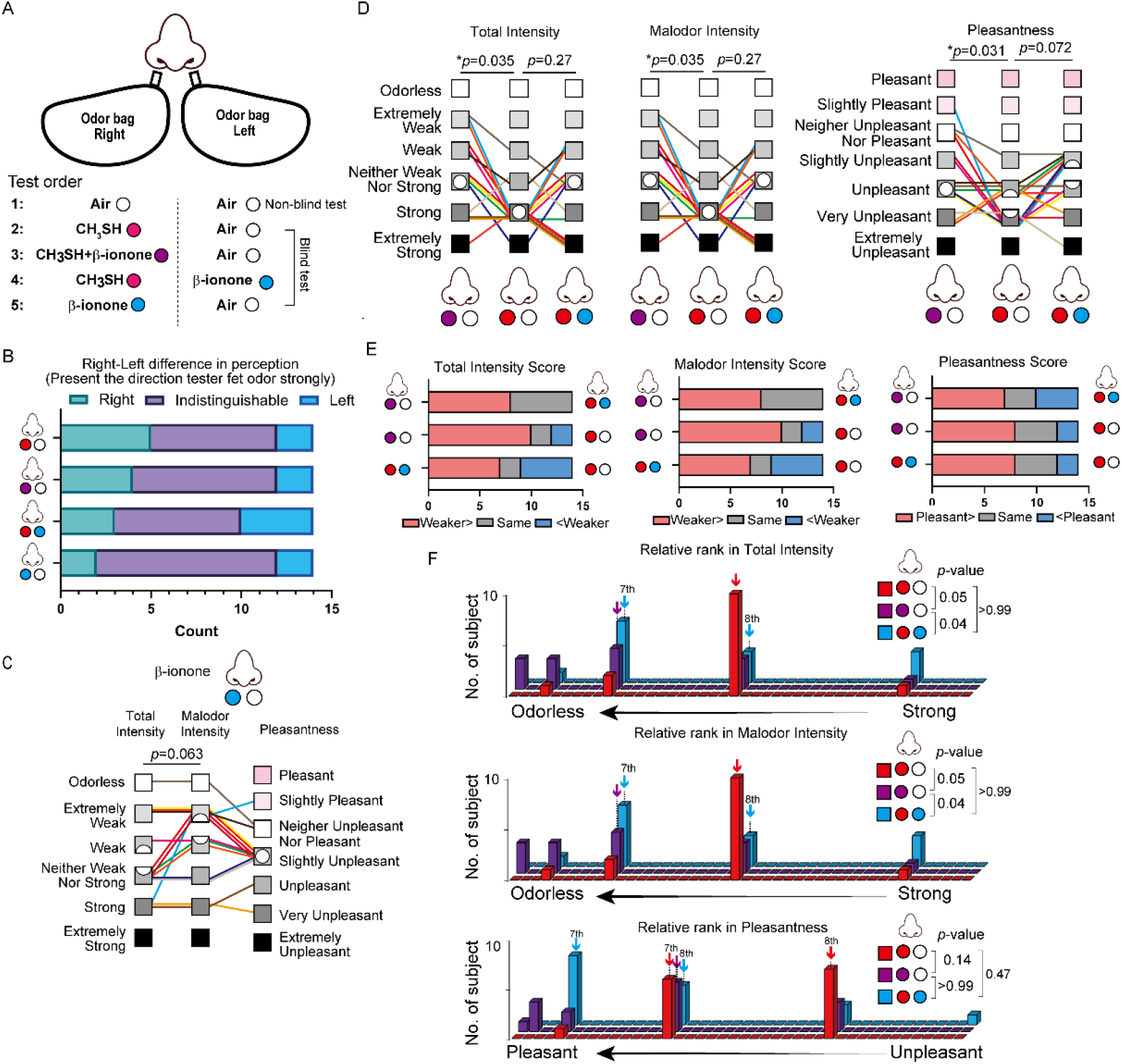
Single nostril sensory evaluation. A) Schematic image of single nostril stimulation in human sensory evaluation test and test odor combinations. B) Test score of whether the malodor comes from the left or right direction in nostril stimulation. Subjects presented the direction they felt strongly. C) Results of evaluation test malodor intensity and total intensity for β-ionone single condition. Comparison test was performed using the non-parametric Wilcoxon test. D) Results of evaluation test for Odor intensity, Malodor intensity and Pleasantness, Center panel: CH_3_SH only, Left panel: mixture gas of CH_3_SH and β-ionone, Right panel: single nostril stimulation by CH_3_SH and β-ionone separately. Each color indicated the score of one subject. Median showed the white circle in each condition. Nonparametric multiple comparisons against mean values were performed using one-way analysis of variance (ANOVA) followed by Dunn’s multiple comparisons test (\**p*<0.05). E) Comparison of which of the stimulus conditions gave the better score. Odor intensity (Left), Malodor intensity (middle) and Pleasantness (Right). F) Relative rank in odor intensity (Upper), malodor intensity (middle) and pleasantness (Lower). Median showed the color arrows in each condition. Both are shown when the 7th and 8th are different. Multiple comparison test was conducted using nonparametric Friedman’s test (\**p*<0.05)

This is consistent with previous reports stating that humans cannot identify the directionality of OSNs when OSNs are stimulated and somatosensory nerves in the nasal cavity are not ^37^. For β-ionone single stimulation at the adjusted concentrations, subjects tended to evaluate that the malodor intensity and pleasantness were neither strong nor weak, and those who felt a strong odor evaluated the malodor intensity higher. (Fig. 4C). Despite being performed in the blind test, all the subjects gave the same score for total odor intensity and malodor intensity for conditions contained methanethiol (Fig. 4D and Supplementary Data). When premixed gas of methanethiol and β-ionone was inhaled from one nostril, total odor intensity, malodor intensity and unpleasantness was reduced with compared to that of each of the components inhaled individually (\**p*=0.035, \**p*=0.035 and *p*=0.031, respectively, Kruskal-Wallis test) (Fig. 4D and 4E). In a striking contrast, when methanethiol and β-ionone were simultaneously inhaled through different nostrils, no significant suppression effect was observed in total odor intensity, malodor intensity and pleasantness (*p*=0.27, *p*=0.27 and *p*=0.072, respectively, Kruskal-Wallis test) (Fig. 4D and 4E). In relative ranking, it is clear that premixed gas showed a significant change in malodor intensity, and that individual stimulation did not (\**p*≤0.05, nonparametric Friedman’s multiple comparison test) (Fig. 4F). Altogether, these results suggest that antagonistic effects of β-ionone on methanethiol-responsive ORs altered odor perception.

## Discussion

OR antagonisms are hypothesized to play an essential role in odor perception, based on previous reports showing widespread antagonistic interactions of odor mixtures at the level of OSNs and ORs in rodents ^15,16,18,38,39^ in conjunction with widespread mixture suppression of odors in humans^24,40^. This study shows that OR2T1 and OR2T11 are activated by methanethiol and antagonized by β-ionone *in vitro*. This corroborates our psychophysics studies showing that β-ionone reduces the intensity and unpleasantness of methanethiol in mixtures. Our results imply that activation of specific OR2T members by certain VSCs induces a characteristic foul odor sensation. Blocking OR2T activation using specific ketone antagonists, such as β-ionone, results in lower odor intensity and unpleasantness of the VSCs, which suggests that OR antagonism plays a prominent role in odor perception Previous studies show that genetic variations of human ORs can cause changes in OR function when tested *in vitro* ^5,41-43^. Loss-of-function variants of ORs are often associated with reduced odor sensation to their ligands, suggesting direct connections between OR activation and odor perception^44^. Consistent with the role of OR2T family members in sulfur odor perception, genetic variation of OR2T6 is associated with liking onions, whose key aroma components are sulfur-containing volatiles ^45^. Additionally, mouse homologs of OR2T family members are among the most significantly activated ORs by sulfur-containing odorants *in vivo* ^46^, supporting the notion that OR2T members are among the most potent ORs against the VSCs.

The binding of sulfur odorants with copper has been studied *in silico* using homology models, which suggested that residues interact with the odorants ^28-30,34,35^ (Extended data Fig. 6). Here, using AlphaFold structural models, we showed that specific residues within the transmembrane domain 5 (C^BW5.43^ and M^BW5.39^) are critical for OR2T1, OR2T6, and OR2T11 binding to hydrogen sulfide and methanethiol complexed with copper. Our models are consistent with those of Haag et al.,^35^ and Vihani et al., ^46^ which noted the potential importance of TM5 in copper-OR complex, notably by a ^5.39^MYxCC^5.43^ motif that is more prevalent in sulfur-activated ORs ^46^. This motif is composed of sulfur-containing amino acids that can coordinate copper and may be how many sulfur-specific ORs bind to sulfur odorants. Our docking simulations also suggest that the β-ionone complex with these ORs occur at this same location, preventing the effective binding of the sulfur agonists.

The use of OR antagonists in evaluating activation/inactivation of specific ORs in odor perception has several advantages over genetic studies. Firstly, a given antagonist can block the activation of multiple related ORs that may have redundant functions. In our study, β-ionone antagonizes OR2T1 and OR2T11 that are activated by methanethiol. If these ORs have similar roles in malodor sensation, it would be difficult to apply genetic methods due to the limited effect size of each genetic variant without assessing a very large number of subjects, as done with the UK biobank study ^47^. Secondly, we can test the same subject to evaluate the effects of an antagonist (and a control) in odor perception, mitigating non-specific effects caused by individuals’ genetic backgrounds and other non-genetic variations. However, a given antagonist for ORs may also act as an agonist for other ORs, creating a potential disadvantage in distinguishing the effects of OR antagonisms from central processing in odor perception. We addressed this challenge by selecting β-ionone, which shows wide variations in perceived odor intensities among subjects ^44,48,49^. We demonstrated that intensity or pleasantness of β-ionone has no correlation with odor suppressive effects against methanethiol. Crucially, the malodor blocking effect was more effective when odorants were presented as a mixture in one nostril than when they were presented separately in each nostril. This strongly supports our hypothesis that β-ionone reduces methanethiol odor intensity and unpleasantness by antagonizing OR2T family members activity.

Our study does not exclude the role of central processing in mixture suppression. In the case of odor masking between 1-propanol and n-amyl, the masking effects were similar when two substances were mixed or presented separately to the two nostrils simultaneously, indicating that the central processing affects odor masking ^23^. In the present study, Iso E Super, which does not potently block OR2T activities, also reduces malodor intensity albeit at lower efficacy, suggesting a role of central processing in masking sulfur odors. Determining the relative roles of antagonistic interactions between odorants and central processing in odor masking in different odor mixtures is a critical step for the future.

Finally, our study highlights a potentially efficient strategy to identify novel deodorants via *in vitro* screening of ORs. This strategy avoids biases, the limited throughput associated with odor adaptation, and the fatigue of human sensory subjects during the initial screening phase. Future studies should determine if this method is fruitful in discovering deodorants for various environmental malodors.

## Acknowledgments

We thank Priyanka Meesa, Emily Xu and Michael Sheyner for reading and editing the manuscript.

## Author Contribution

Y.F. conceived and designed the project. Y.F., M.A. and H.S. performed the ligand assay. R.E. and T.T. conducted sensory evaluation test. C.D.M. performed ligand docking and binding analysis, Y.F., H.M. and M.Y. carried out the analysis and wrote the paper with inputs from all authors. Y.F., H.M. and Y.M. supervised the project.

## Correspondence

Correspondence should be addressed to Y.F., H.M. and M.Y..

## Methods

### DNA and vector preparation

Open reading frames of human OR genes were subcloned into pCI (Promega, WI, USA) with a Rho-tag (the sequence encoding the first 20 amino acids of rhodopsin) at the N terminal. To generate mutants of ORs, DNA fragments of OR genes were amplified by PrimeStar MAX polymerase (Takara bio, Shiga, Japan). The fragments were mixed and amplified by PCR reaction to obtain full sequences. The plasmid for the expression of human ORs, RTP1S^50,51^, and pGlosensor F-22 (Promega) were amplified and purified by Nucleospin plasmid TF grade (Takara bio, Shiga, Japan). All plasmid sequences were verified using Sanger sequencing (3100 Genetic Analyzer, Applied Biosystems).

### Cell culture

HEK293T and Hana 3A cells ^52^ were grown in Minimal Essential Medium (MEM) containing 10% FBS (vol/vol) with penicillin-streptomycin and amphotericin B. Hana 3A cells were authenticated using polymorphic short tandem repeat (STR) at the Duke DNA Analysis Facility using GenePrint 10 (Promega) and shown to share profiles with the reference (ATCC). All cell lines were incubated at 37°C, saturating humidity, and 5% CO2. No mycoplasma infection was detected in all cell cultures.

### Vapor Glosensor assay

In the volatile sulfur detection test, Vapor Glosensor cAMP Assay (Promega) was used to measure the changes in cAMP levels caused by receptor activation upon ligand binding^11^. Hana3A cells were plated on poly-D-Lysine coated 96-well plates. 18-24 hours after plating, cells were transfected with 80 ng/well of plasmids encoding ORs, 5 ng/well of RTP1S ^52^, and 10 ng/well of Glosensor plasmid (Promega). 18-24 hours later, the medium was replaced with 25 μL of HBSS (Gibco) containing 10 mM HEPES and 1 mM Glucose, followed by 25 μL of the HBSS containing GloSensor cAMP Reagent (Promega). Plates were kept in a dark place at room temperature for two hours to equilibrate cells with the reagent. The test plate was inserted into the plate reader. The luminescence derived from basal activity in each ORs was measured. Before odor stimulation of the cells expressing individual ORs on testing 96 well plate by odorants, a 96 well plate put into the 5L PET film sampling bag (Flek-Sampler, Omi odor air service Co., Shiga, Japan) with a small fan. After 5L of pure air gas was inserted into the bag containing the assay plate, H_2_S or CH_3_SH gas was added into same bag to make the desired gas phase concentration. Based on a previous report that an odor response of the heterologous cells expressing ORs is more than 1000 times weaker than that of olfactory sensory neuron cells^12^, the stimulation concentration for 1st screening (7 ppm for CH_3_SH and 41 ppm for H_2_S) was set to 10,000 times the human olfactory threshold concentration (0.00007 ppm for CH_3_SH and 0.00041 ppm for H_2_S)^53^. Concentration of VSCs in sampling bag was measured with the Gas detector tube system and detector tubes (No.70L for CH_3_SH and No.4LL for H_2_S) (GASTEC Corporation, Kanagawa, Japan). Then the assay plate was incubated in the bag for 10 minutes. Immediately, the test plate was inserted in the plate reader GloMAX discover (Promega) to measure the luminescence by VSC stimulation. The luminescence in each well was measured. Response was evaluated by the fold change in luminescence value before and after odor stimulation.

When the screening and evaluating antagonist was conducted, before putting the assay plate into the sampling bag, tested fragrance solutions were put into each well on the assay plate. When evaluating the response to each fragrance against ORs, all luminescence values were divided by the value obtained from the cells transfected with the empty vector at the same cycle. Multiple comparisons were performed using one-way analysis of variance (ANOVA) followed by Dunnett’s test with the target group set to control condition.

### Docking

Structures were downloaded from the Alfaphold 2 database (https://alphafold.ebi.ac.uk/, downloaded 02/17/2022) ^54^. The most variable parts of the receptor were mainly located at the Nter (Fig S12). OR2T1 possesses an amino acid abnormally longer Nter loop than mammals OR (Fig S12 and S13) that is modelled with low confidence by Alphafold. In consequence, we decided to truncate the Nter parts of the models as indicated in Fig S13. The structures of OR2T1, OR2T6, and OR2T11 are very close as the Root Mean Square Deviation (RMSD) between the structures are very low (Fig S12). Odorant (hydrogen sulfide, methanethiol and β-ionone) and copper SDF files were downloaded from Pubchem ^55^ and converted in pdb with Open Babel 2.3.2 on PyRx 0.8 ^56^ and in pdbqt with Autodock Tools 1.5.7 ^57^. The truncated Alfaphold models were converted in pdbqt files and the grid for docking ligands and copper was generated with Autodock Tools (Fig S14). Docking was realized with Vina 1.2.0 ^58,59^. On a rigid receptor, 20 poses were generated with an energy range of 15 kcal/mol and an exhaustiveness of 10. Results were analyzed on Autodock Tools, and visualized on VMD 1.9.3^60^ and UCSF Chimera 1.15^61^.

### Sensory Evaluation test

All procedures involving human subjects were approved by the ethics committee of ST Corporation. All subjects gave informed consent to participate. A filter paper placed in a beaker was impregnated with 1g of undiluted solution of fragrances. The beaker was placed in a 10L sampling bag filled with non-odor pure air gas filtered through silica and activated charcoal.

The scented air was adjusted by allowing the bag to stand overnight at room temperature. A new 3L sampling bag was willed with pure air gas. Then, 200ml of the scented air and 0.4 mL of 2% CH_3_SH gas (final concentration of 0.3 ppm) were injected into the bag. In the same process each fragrance and CH_3_SH gas alone were adjusted. Each odor bag was prepared for the whole nose test and single-nostril test (Extended data Fig. 10). Subjects evaluated the total intensity, malodor intensity and pleasantness against each condition. The example of answer sheet shown in Extended data Fig. 10 was used for the evaluation. In order to clarify the evaluation criteria, a non-blind test was conducted for Methanthiol alone and the other test in the blind condition. In the test for β-ionone and Iso E Super, panelists (n=20) were divided into two, and the order of the scents was changed for each. No grouping is performed for sensory evaluation using other fragrances. The panelists were men and women in their 20s and 30s.

Detailed data of each panelist was also shown in Supplementary Data. The panelists evaluated the degree of pleasant/unpleasant, the intensity of malodor, and the intensity of total odor.

Single nostril evaluation tests were performed in non-blind condition against the control bag containing only pure air and the other test in the blind condition. Panelists were instructed to sniff their hands and arms and reset their noses between samples. Nonparametric multiple comparisons were performed using one-way analysis of variance (ANOVA) following Dunn’s or Friedman’s multiple comparisons test using the GraphPad Prism. Correlation values were also calculated by nonparametric Spearman correction test.

## Statistical analysis

Multiple comparisons were performed using one-way analysis of variance (ANOVA) using the GraphPad Prism. Analysis methods for each data were also described in each methods and figure legends.

## Data availability

All relevant data are available within the manuscript and its supplementary information or from the authors upon reasonable request.

## Supplementary Figures

**Extended data Figure 1.**
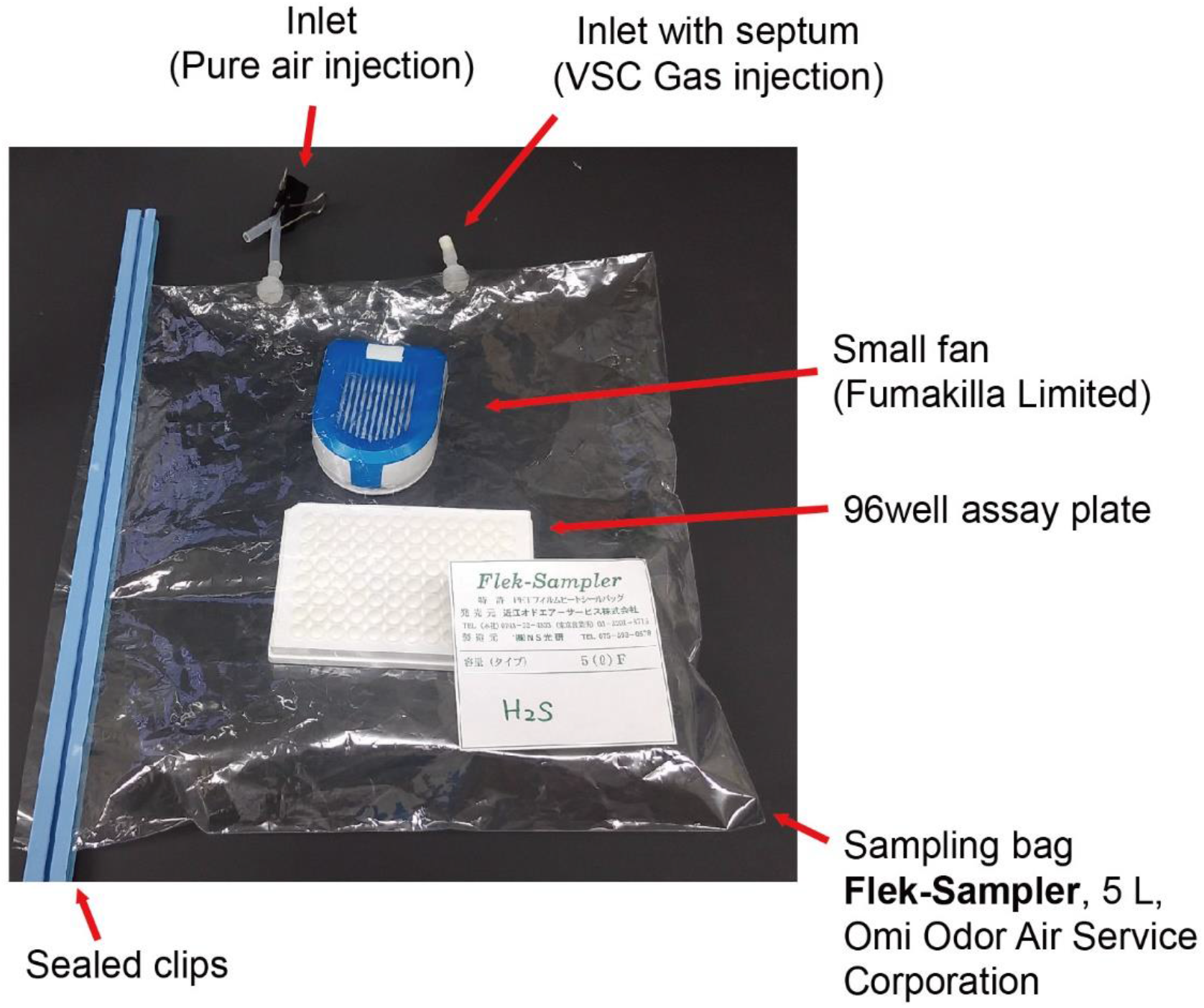
Photo image of the vapor stimulation assay with the cell culturing 96 well assay plate and the small fan in the sampling bag.

**Extended data Figure 2.**
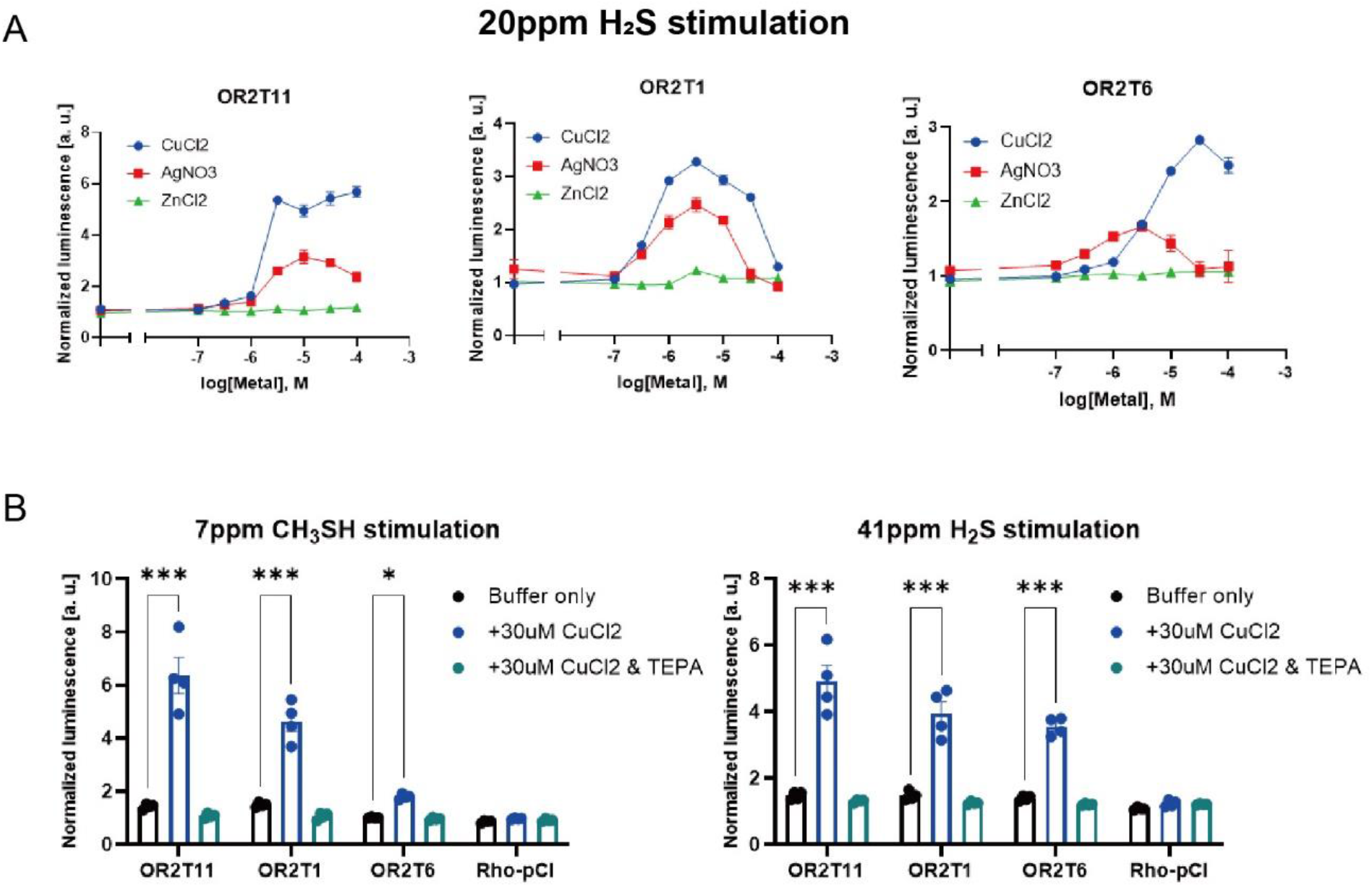
VSC binding on odorant receptors via copper ions. A metal ion is essential for CH3SH/H_2_S responding ORs. A) Dose-response curves of 3 ORs against increasing concentrations of metals with the concentration of vapor H_2_S constant at 20 ppm. B) Cu^2+^ Cheater TEPA diminished the response of ORs against both of H_2_S and CH_3_SH. The y-axis indicates normalized response± s.e.m (n=3). Multiple comparisons were performed using one-way analysis of variance (ANOVA) followed by Dunnett’s multiple comparison test (\**p*<0.05. \*\**p*<0.01, \*\*\**p*<0.001).

**Extended data Figure 3.**
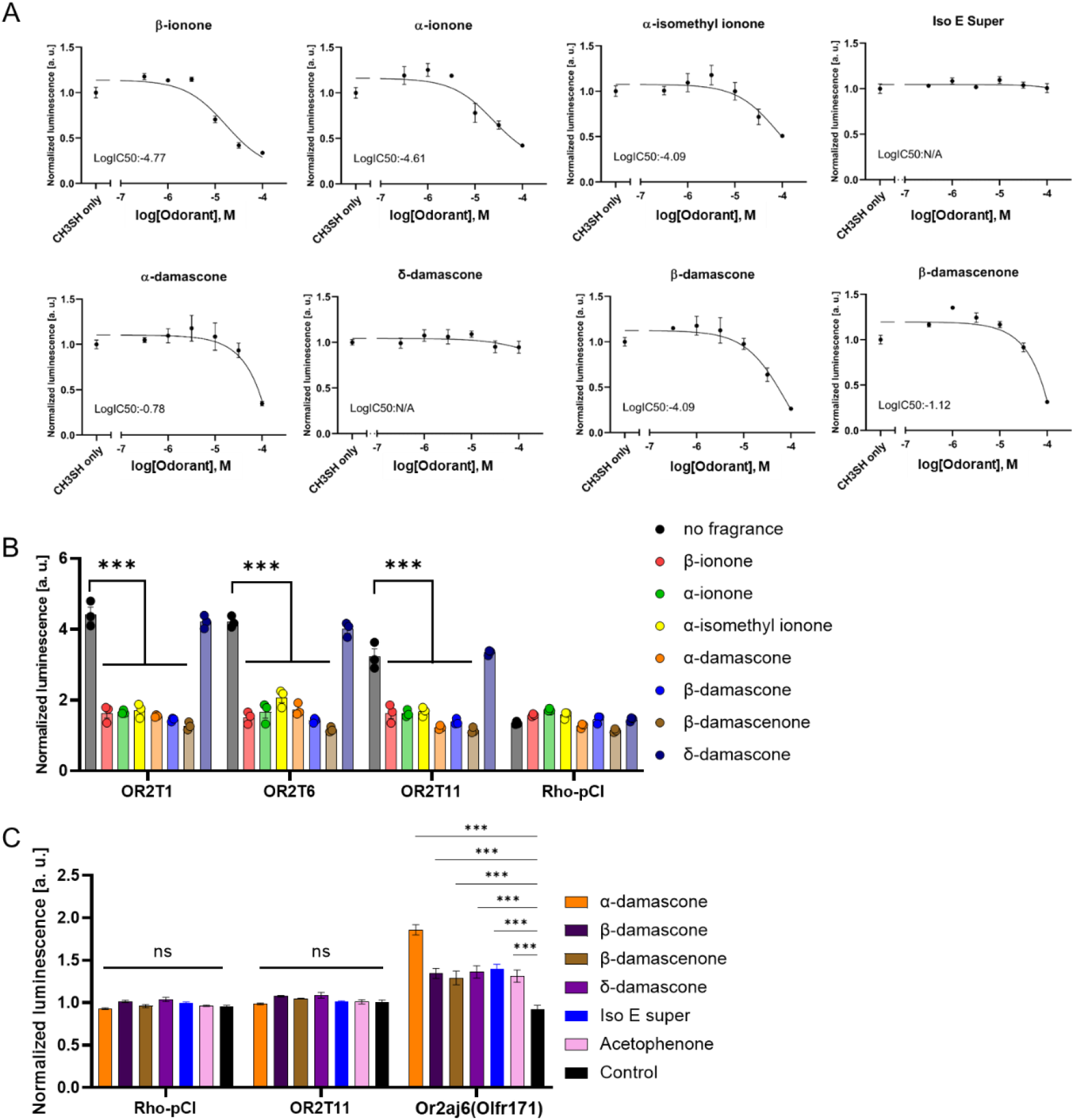
Dose response analysis of candidate antagonists. A) Dose-response curves of OR2T11 against increasing concentrations of ionones and damascones, with the vapor concentration of CH_3_SH held constant at 7 ppm. IC50 value was calculated using Graph pad Prism software. B) Antagonistic effect of damascone & ionone analogs. 41 ppm H_2_S stimulation on human OR2T1, OR2T6 and OR2T11 were masked by 100 µM fragrance compounds. Multiple comparisons were performed using one-way analysis of variance (ANOVA) followed by Dunnett’s multiple comparison test (\*\*\**p*<0.001). C) Inhibitor odorants did not cause adverse effects on the assay system. OR2T11 and mouse Or2aj6 (Olfr171) (damascones responding receptor) expressing cells were stimulated by 100µM odorants and Glosensor buffer without any odorant as a negative control. Error bars indicate s.e.m (n=3) Multiple comparisons were performed using one-way ANOVA followed by Dunnett’s test (\*\*\**p*<0.001).

**Extended data Figure 4.**
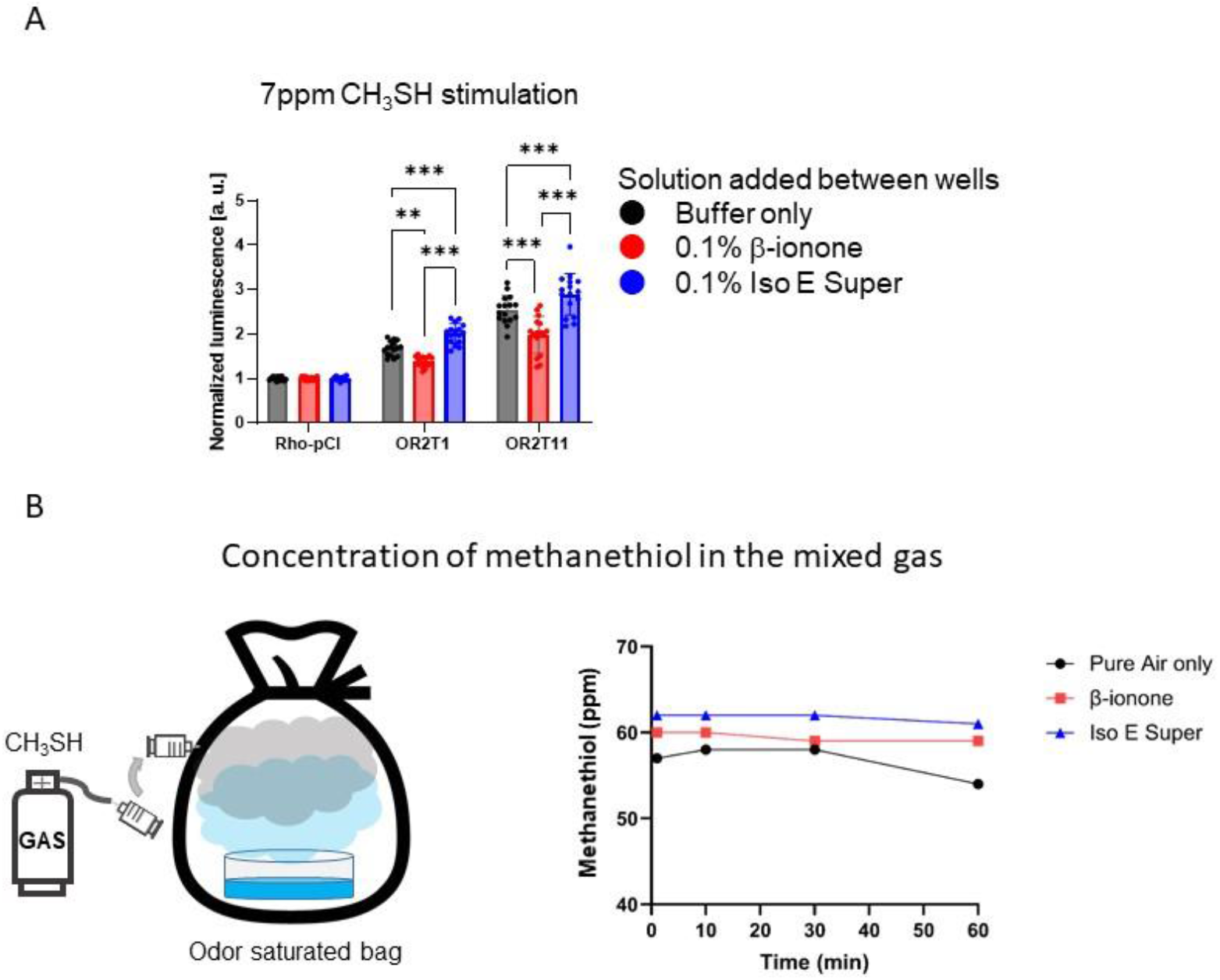
Inhibitory effects of β-ionone in the vapor phase. A) Inhibitory effect on vapor phase mixing β-ionone or Iso E super. Soon after adding 25 µL of β-ionone or Iso E Super solution between the wells of the 96-well plate, OR expressing cells was quickly stimulated by 7ppm CH_3_SH gas for 10 min. The normalized luminescence value indicates the S/N ratio before and after stimulation of CH_3_SH. Multiple comparisons were performed using one-way analysis of variance (ANOVA) followed by Dunnett’s Test (\**p*<0.05. \*\**p*<0.01) B) Concentration of methanethiol in the sampling bag containing β-ionone or Iso E Super. methanethiol gas was injected into the saturated gas phase of β-ionone or Iso E Super in the sampling bag. Concentration of methanthiol in sampling bag was measured with the Gas detector tube system and detector tube No.70L for CH_3_SH detection.

**Extended data Figure 5.**
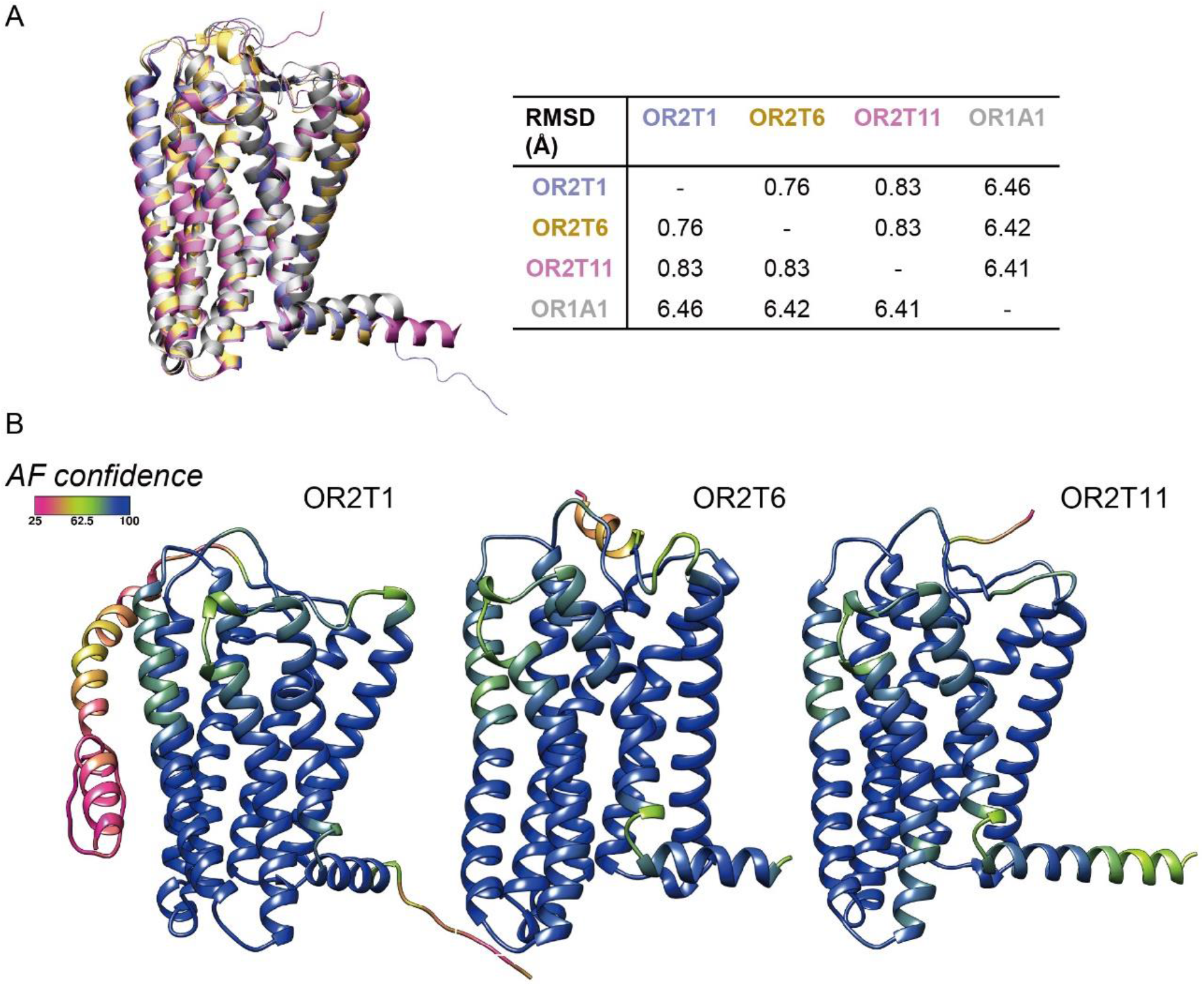
3D model of volatile sulfur responding ORs. A) Superimposition of the Alphafold models of OR2T1, OR2T6 and OR2T11. RMSD between the backbone of the structures is shown in a table. B) Alphafold confidence score projected on the Alphafold models of OR2T1, OR2T6 and OR2T11.

**Extended data Figure 6.**
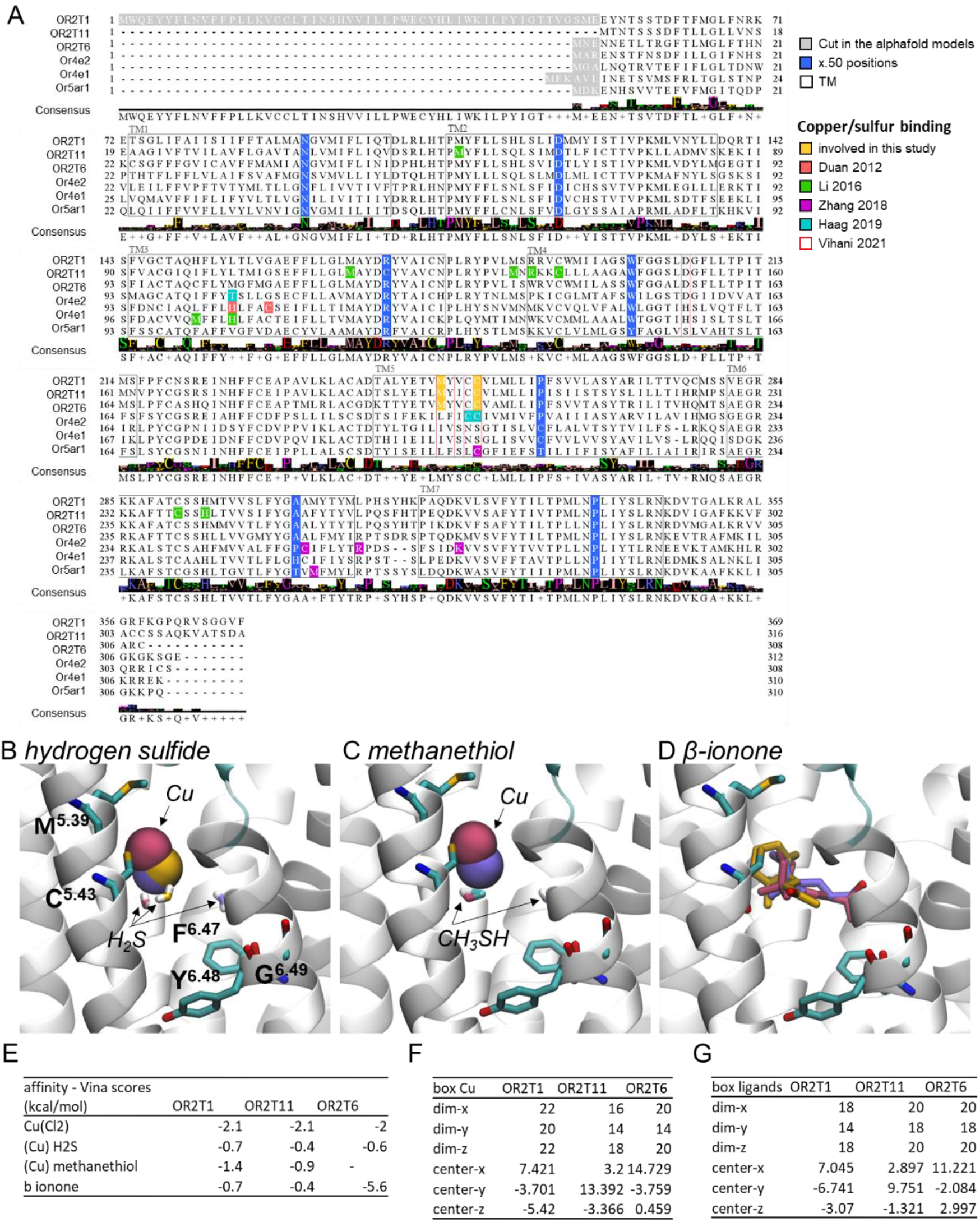
Simulation analysis of inhibitory effects by β-ionone. A) Alignment of the sulfur responding OR studied in the literature. X.50 positions, TM domains as well as amino acids studied here and previously identified are highlighted. B-D) Docking results for H_2_S (B), methanethiol (C) and β-ionone (D) for OR2T1 (purple), OR2T6 (yellow) and OR2T11 (pink). Copper is represented in Van der Waals volume colored by OR while the ligand is represented in licorice with sulfur (H_2_S and methanethiol) or carbon (β-ionone) atom colored by OR. E) Docking results by affinity represented by the Vina score. F) Box dimension and placement for copper docking. G) Box dimension and placement for ligand docking.

**Extended data Figure 7.**
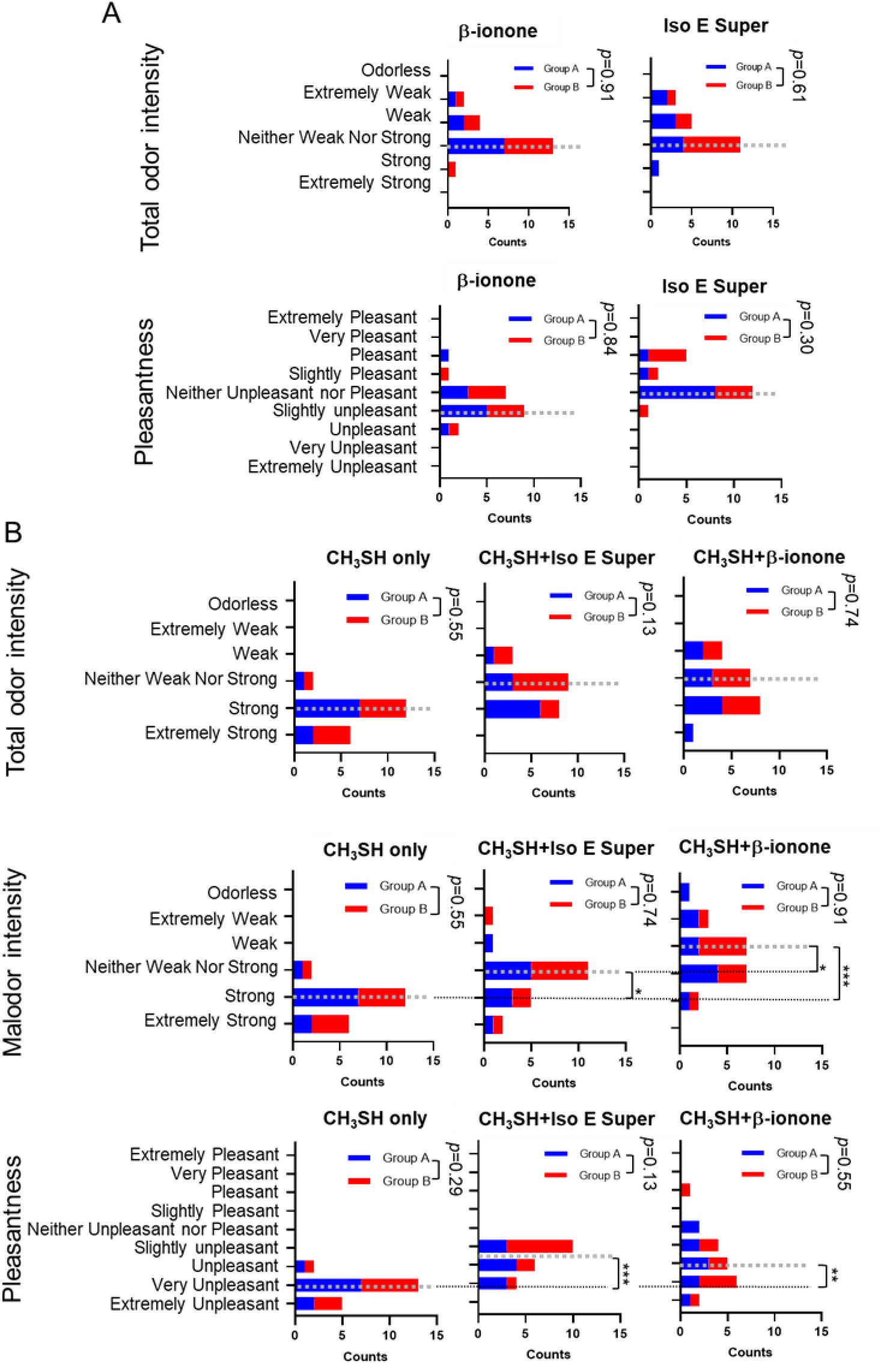
Sensory evaluation test. A) Score for β-ionone or Iso E Super. Red: Group A (n=10), Blue: Group B (n=10), Dot line: Median value. B) Score for methanethiol only, methanethiol and Iso E Super, and methyl methanethiol and β-ionone. Red: Group A (n=10), Blue: Group B (n=10), Dot line: Median value. The significance analysis between Group A and B was performed using Mann-Whitney test. Nonparametric multiple comparisons among mixed gas were performed using one-way analysis of variance (ANOVA) followed by Dunn’s multiple comparisons test (\**p*<0.05. \*\**p*<0.01, \*\*\**p*<0.001).

**Extended data Figure 8.**
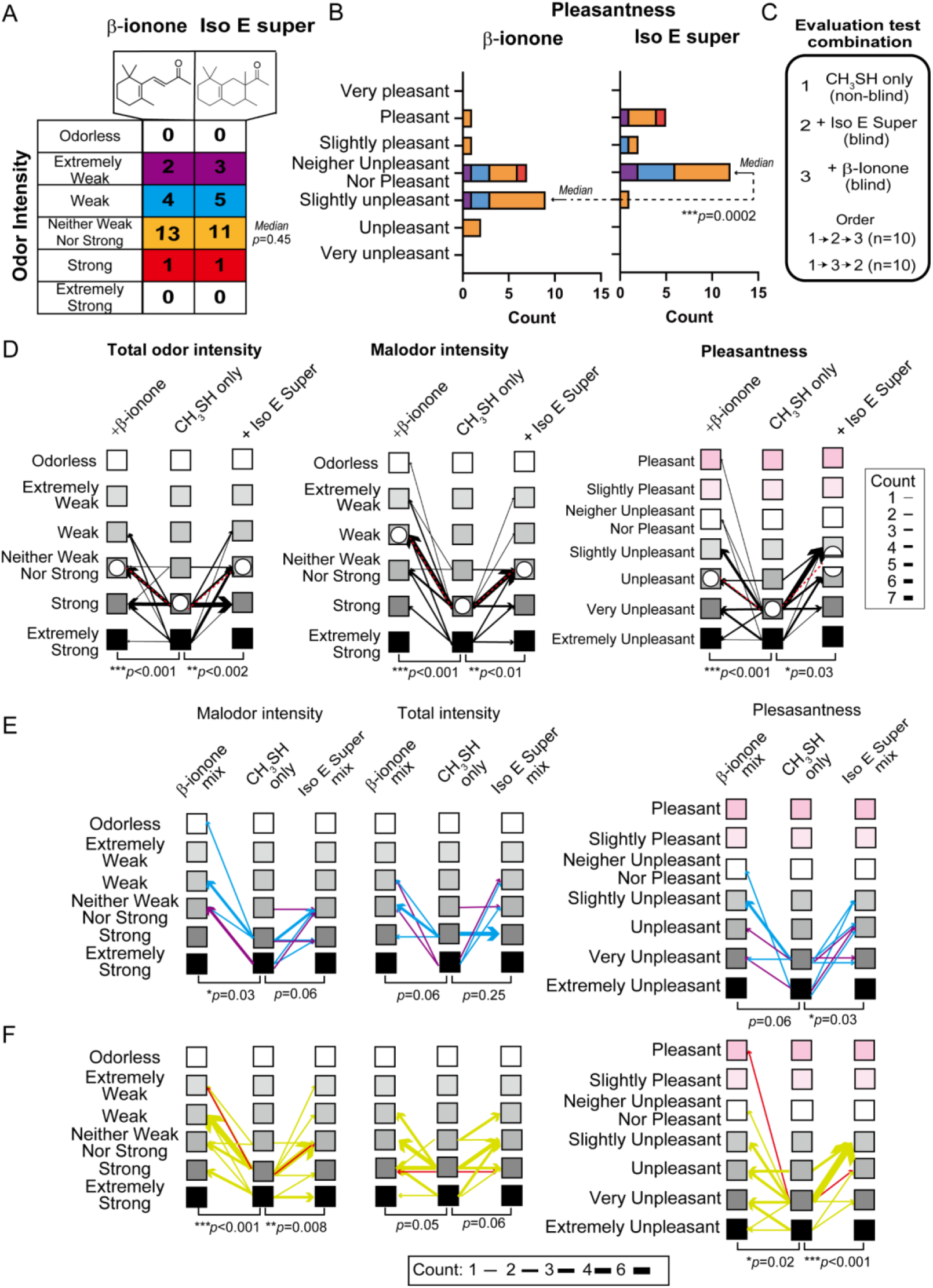
Human Sensory evaluation test against CH_3_SH gas containing β-ionone (an effective antagonist) or Iso E Super. A) Odor intensity value of β-ionone and Iso E Super evaluated by 20 subjects. B) Pleasantness value of β-ionone and Iso E Super. Color matched the odor intensity score of each subject representing in Extended Fig. 8A. Comparison test was performed using the non-parametric Wilcoxon test Test (***p<0.001). C) Test odor combination and order in sensory evaluation test. D) Results of evaluation test Total odor intensity Malodor intensity and Pleasantness. Line thickness: The number of subjects who made the same evaluation transition in mixture gas. Median showed the white circle in each condition. E and F) Results of evaluation test Total odor intensity Malodor intensity and Pleasantness when classified into Odor intensity scores of β-ionone or Iso E super. Color matched the odor intensity score of each subject representing in Extended Fig 8A (E: Extremely weak and Weak, F: Neither weak nor strong and Strong). Line thickness: The number of subjects who made the same evaluation transition in mixture gas. Comparison test was performed using the non-parametric Wilcoxon test Test.

**Extended data Figure 9.**
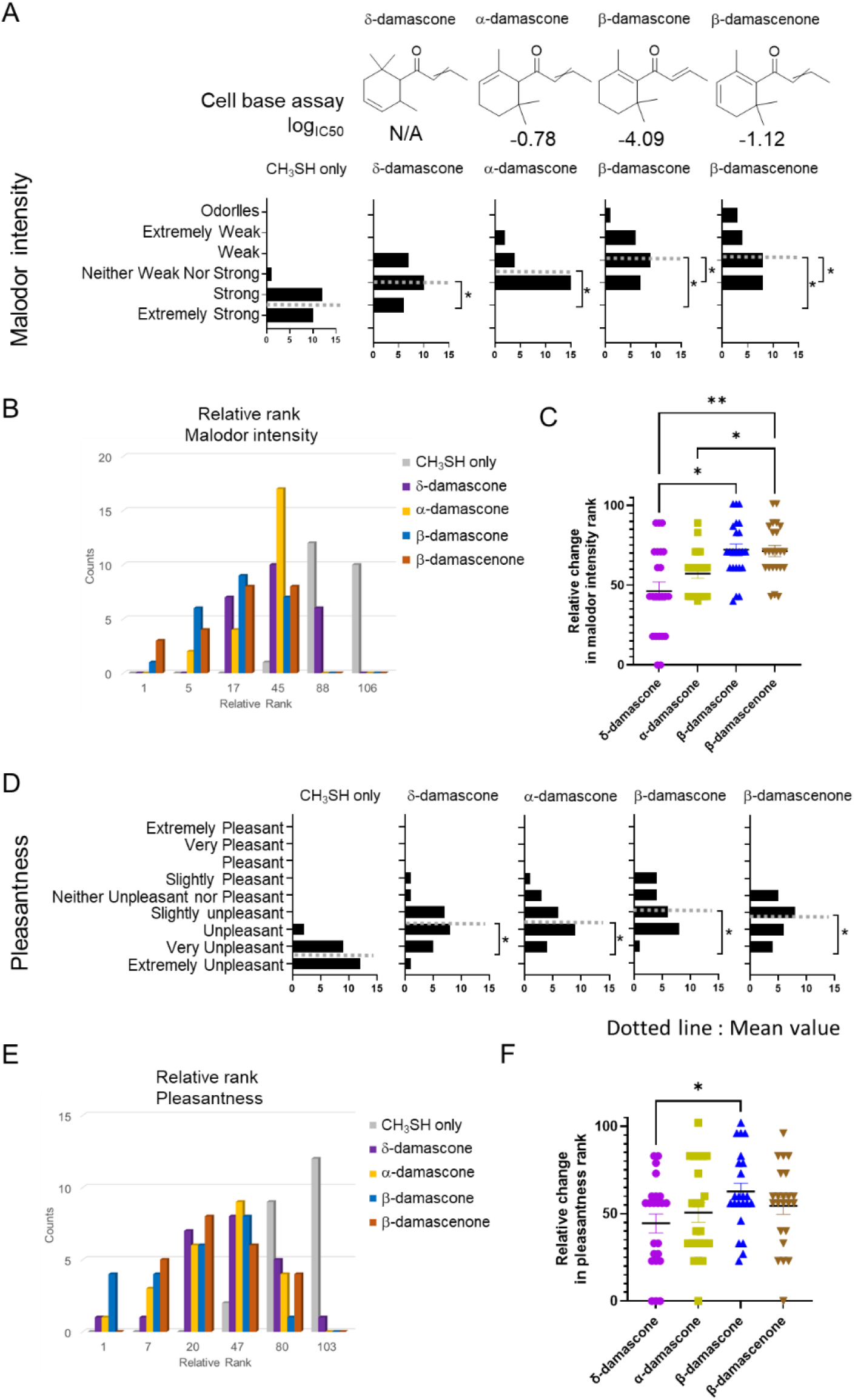
Damascones changes odor perception against VSC. A and D) Sensory evaluation score for methanethiol only and mixture gas with damascones. A: Odor intensity, D: Pleasantness, Gray dot line: Mean value. Nonparametric multiple comparisons were performed using one-way analysis of variance (ANOVA) followed by Dunn’s multiple comparisons test (\**p*<0.05). B and E) Relative rank in malodor intensity and pleasantness, C and F) Relative rank change in malodor intensity (C) and pleasantness (F) Multiple comparison test was conducted using nonparametric Friedman’s test (\**p*<0.05, \*\**p*<0.01).

**Extended data Figure 10.**
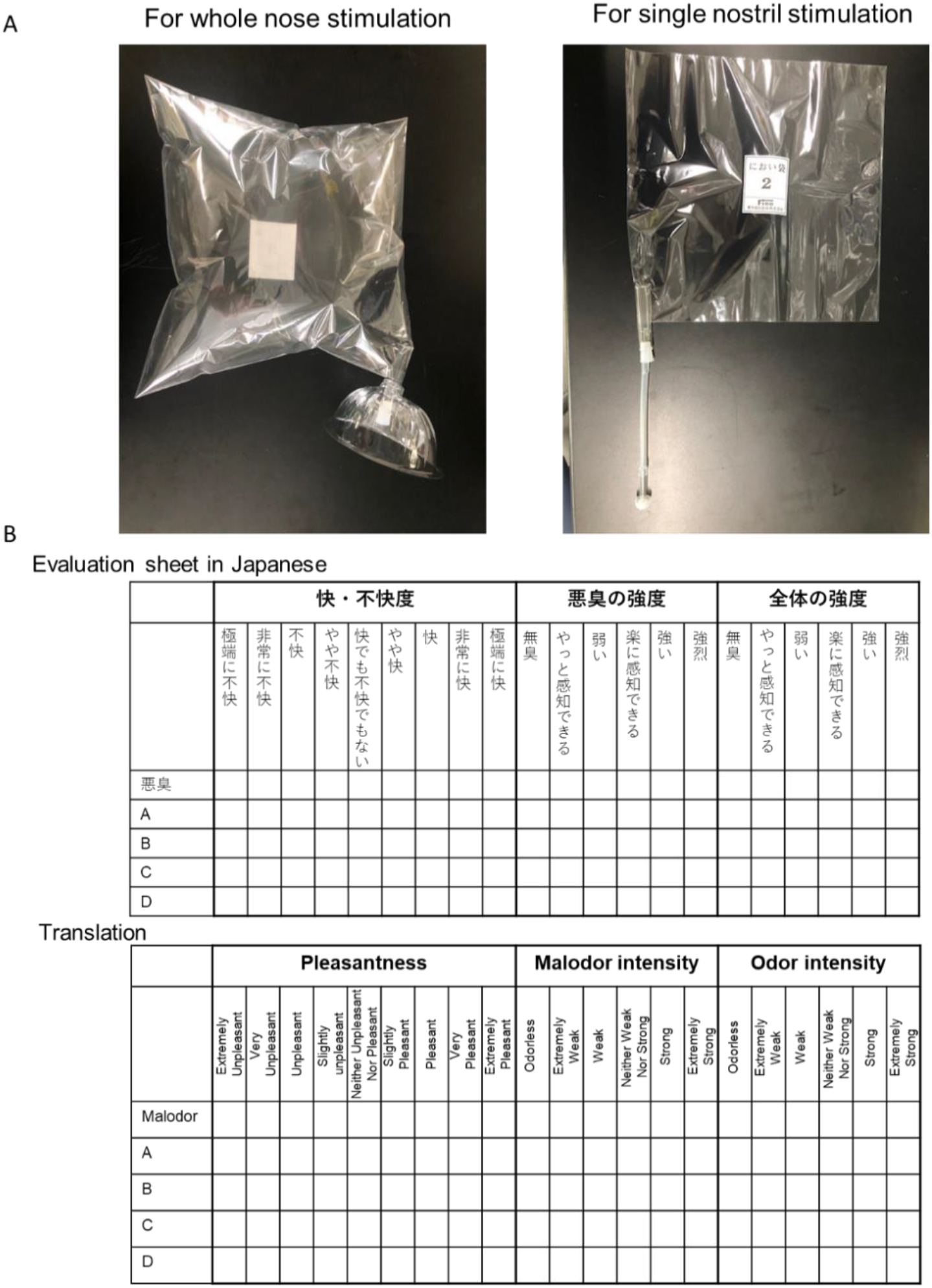
Equipment for sensory evaluation test. Left) Photographic image of the sampling bags used in sensory evaluation test. Upper) for whole nose stimulation (Results were shown in Figure 3), Lower) for single nostril stimulation test (Results were shown in Figure 4). Each has a suction port of a suitable size. Right) Score sheet used in sensory evaluation tests. Upper: original score sheet written in Japanese. Lower: English translation version of original.

## References

1 Buck, L. & Axel, R. A novel multigene family may encode odorant receptors: a molecular basis for odor recognition. Cell 65, 175–187, doi:10.1016/0092-8674(91)90418-x (1991).

2 Niimura, Y., Matsui, A. & Touhara, K. Extreme expansion of the olfactory receptor gene repertoire in African elephants and evolutionary dynamics of orthologous gene groups in 13 placental mammals. Genome Res 24, 1485–1496, doi:10.1101/gr.169532.113 (2014).

3 Adipietro, K. A., Mainland, J. D. & Matsunami, H. Functional evolution of mammalian odorant receptors. PLoS Genet 8, e1002821, doi:10.1371/journal.pgen.1002821 (2012).

4 Malnic, B., Godfrey, P. A. & Buck, L. B. The human olfactory receptor gene family. Proc Natl Acad Sci U S A 101, 2584–2589, doi:10.1073/pnas.0307882100 (2004).

5 Trimmer, C. et al. Genetic variation across the human olfactory receptor repertoire alters odor perception. Proc Natl Acad Sci U S A 116, 9475–9480, doi:10.1073/pnas.1804106115 (2019).

6 Serizawa, S., Miyamichi, K. & Sakano, H. One neuron-one receptor rule in the mouse olfactory system. Trends Genet 20, 648–653, doi:10.1016/j.tig.2004.09.006 (2004).

7 Ferreira, T. et al. Silencing of odorant receptor genes by G protein betagamma signaling ensures the expression of one odorant receptor per olfactory sensory neuron. Neuron 81, 847–859, doi:10.1016/j.neuron.2014.01.001 (2014).

8 Saito, H., Chi, Q., Zhuang, H., Matsunami, H. & Mainland, J. D. Odor coding by a Mammalian receptor repertoire. Sci Signal 2, ra9, doi:10.1126/scisignal.2000016 (2009).

9 Nara, K., Saraiva, L. R., Ye, X. & Buck, L. B. A large-scale analysis of odor coding in the olfactory epithelium. J Neurosci 31, 9179–9191, doi:10.1523/JNEUROSCI.1282-11.2011 (2011).

10 Saraiva, L. R. et al. Combinatorial effects of odorants on mouse behavior. Proc Natl Acad Sci U S A 113, E3300–3306, doi:10.1073/pnas.1605973113 (2016).

11 Kida, H. et al. Vapor detection and discrimination with a panel of odorant receptors. Nat Commun 9, 4556, doi:10.1038/s41467-018-06806-w (2018).

12 Oka, Y., Omura, M., Kataoka, H. & Touhara, K. Olfactory receptor antagonism between odorants. EMBO J 23, 120–126, doi:10.1038/sj.emboj.7600032 (2004).

13 Liu, M. T. et al. Carbon chain shape selectivity by the mouse olfactory receptor OR-I7. Org Biomol Chem 16, 2541–2548, doi:10.1039/C8OB00205C (2018).

14 Singh, V., Murphy, N. R., Balasubramanian, V. & Mainland, J. D. Competitive binding predicts nonlinear responses of olfactory receptors to complex mixtures. Proc Natl Acad Sci U S A 116, 9598–9603, doi:10.1073/pnas.1813230116 (2019).

15 Xu, L. et al. Widespread receptor-driven modulation in peripheral olfactory coding. Science 368, doi:10.1126/science.aaz5390 (2020).

16 Inagaki, S., Iwata, R., Iwamoto, M. & Imai, T. Widespread Inhibition, Antagonism, and Synergy in Mouse Olfactory Sensory Neurons In Vivo. Cell Rep 31, 107814, doi:10.1016/j.celrep.2020.107814 (2020).

17 Pfister, P. et al. Odorant Receptor Inhibition Is Fundamental to Odor Encoding. Curr Biol 30, 2574–2587 e2576, doi:10.1016/j.cub.2020.04.086 (2020).

18 de March, C. A. et al. Modulation of the combinatorial code of odorant receptor response patterns in odorant mixtures. Mol Cell Neurosci 104, 103469, doi:10.1016/j.mcn.2020.103469 (2020).

19 Reisert, J. Origin of basal activity in mammalian olfactory receptor neurons. J Gen Physiol 136, 529–540, doi:10.1085/jgp.201010528 (2010).

20 Brodin, M., Laska, M. & Olsson, M. J. Odor interaction between Bourgeonal and its antagonist undecanal. Chem Senses 34, 625–630, doi:10.1093/chemse/bjp044 (2009).

21 Spehr, M. et al. Dual capacity of a human olfactory receptor. Curr Biol 14, R832–833, doi:10.1016/j.cub.2004.09.034 (2004).

22 Burger, J. L., Jeerage, K. M. & Bruno, T. J. Direct nuclear magnetic resonance observation of odorant binding to mouse odorant receptor MOR244-3. Anal Biochem 502, 64–72, doi:10.1016/j.ab.2016.03.006 (2016).

23 Cain, W. S. Odor Intensity - Mixtures and Masking. Chem Sens Flav 1, 339–352, doi:DOI 10.1093/chemse/1.3.339 (1975).

24 Thomas-Danguin, T. et al. The perception of odor objects in everyday life: a review on the processing of odor mixtures. Frontiers in psychology 5, 504, doi:10.3389/fpsyg.2014.00504 (2014).

25 Kurahashi, T., Lowe, G. & Gold, G. H. Suppression of odorant responses by odorants in olfactory receptor cells. Science 265, 118–120, doi:10.1126/science.8016645 (1994).

26 Wallrabenstein, I., Singer, M., Panten, J., Hatt, H. & Gisselmann, G. Timberol(R) Inhibits TAAR5-Mediated Responses to Trimethylamine and Influences the Olfactory Threshold in Humans. PLoS One 10, e0144704, doi:10.1371/journal.pone.0144704 (2015).

27 de March, C. A., Fukutani, Y., Vihani, A., Kida, H. & Matsunami, H. Real-time In Vitro Monitoring of Odorant Receptor Activation by an Odorant in the Vapor Phase. J Vis Exp, doi:10.3791/59446 (2019).

28 Li, S. et al. Smelling Sulfur: Copper and Silver Regulate the Response of Human Odorant Receptor OR2T11 to Low-Molecular-Weight Thiols. J Am Chem Soc 138, 13281–13288, doi:10.1021/jacs.6b06983 (2016).

29 Block, E., Batista, V. S., Matsunami, H., Zhuang, H. & Ahmed, L. The role of metals in mammalian olfaction of low molecular weight organosulfur compounds. Nat Prod Rep 34, 529–557, doi:10.1039/c7np00016b (2017).

30 Zhang, R. et al. A Multispecific Investigation of the Metal Effect in Mammalian Odorant Receptors for Sulfur-Containing Compounds. Chemical senses 43, 357–366, doi:10.1093/chemse/bjy022 (2018).

31 Gautschi, M., Bajgrowicz, J. A. & Kraft, P. Fragrance chemistry - Milestones and perspectives. Chimia 55, 379–387 (2001).

32 Armanino, N. et al. What’s Hot, What’s Not: The Trends of the Past 20 Years in the Chemistry of Odorants. Angew Chem Int Ed Engl 59, 16310–16344, doi:10.1002/anie.202005719 (2020).

33 Jumper, J. et al. Highly accurate protein structure prediction with AlphaFold. Nature 596, 583–589, doi:10.1038/s41586-021-03819-2 (2021).

34 Duan, X. et al. Crucial role of copper in detection of metal-coordinating odorants. Proc Natl Acad Sci U S A 109, 3492–3497, doi:10.1073/pnas.1111297109 (2012).

35 Haag, F. et al. Copper-mediated thiol potentiation and mutagenesis-guided modeling suggest a highly conserved copper-binding motif in human OR2M3. Cell Mol Life Sci, doi:10.1007/s00018-019-03279-y (2019).

36 de March, C. A. et al. Conserved Residues Control Activation of Mammalian G Protein-Coupled Odorant Receptors. J Am Chem Soc 137, 8611–8616, doi:10.1021/jacs.5b04659 (2015).

37 Wu, Y., Chen, K., Ye, Y., Zhang, T. & Zhou, W. Humans navigate with stereo olfaction. Proc Natl Acad Sci U S A 117, 16065–16071, doi:10.1073/pnas.2004642117 (2020).

38 Reddy, G., Zak, J. D., Vergassola, M. & Murthy, V. N. Antagonism in olfactory receptor neurons and its implications for the perception of odor mixtures. Elife 7, doi:10.7554/eLife.34958 (2018).

39 Zak, J. D., Reddy, G., Vergassola, M. & Murthy, V. N. Antagonistic odor interactions in olfactory sensory neurons are widespread in freely breathing mice. Nat Commun 11, 3350, doi:10.1038/s41467-020-17124-5 (2020).

40 Ishii, A. et al. Synergy and masking in odor mixtures: an electrophysiological study of orthonasal vs. retronasal perception. Chem Senses 33, 553–561, doi:10.1093/chemse/bjn022 (2008).

41 Keller, A., Zhuang, H., Chi, Q., Vosshall, L. B. & Matsunami, H. Genetic variation in a human odorant receptor alters odour perception. Nature 449, 468–472, doi:10.1038/nature06162 (2007).

42 Mainland, J. D. et al. The missense of smell: functional variability in the human odorant receptor repertoire. Nat Neurosci 17, 114–120, doi:10.1038/nn.3598 (2014).

43 McRae, J. F. et al. Genetic variation in the odorant receptor OR2J3 is associated with the ability to detect the “grassy” smelling odor, cis-3-hexen-1-ol. Chem Senses 37, 585–593, doi:10.1093/chemse/bjs049 (2012).

44 Jaeger, S. R. et al. A Mendelian trait for olfactory sensitivity affects odor experience and food selection. Curr Biol 23, 1601–1605, doi:10.1016/j.cub.2013.07.030 (2013).

45 May-Wilson, S. et al. Large-scale genome-wide association study of food liking reveals genetic determinants and genetic correlations with distinct neurophysiological traits. bioRxiv, 2021.2007.2028.454120, doi:10.1101/2021.07.28.454120 (2021).

46 Vihani, A. et al. Encoding of odors by mammalian olfactory receptors. bioRxiv, 2021.2012.2027.474279, doi:10.1101/2021.12.27.474279 (2021).

47 Sanchez-Roige, S., Gray, J. C., MacKillop, J., Chen, C. H. & Palmer, A. A. The genetics of human personality. Genes Brain Behav 17, e12439, doi:10.1111/gbb.12439 (2018).

48 Plotto, A., Barnes, K. W. & Goodner, K. L. Specific Anosmia Observed for β-Ionone, but not for α-Ionone: Significance for Flavor Research. J. Food Sci. 71, S401–S406 (2006).

49 McRae, J. F. et al. Identification of regions associated with variation in sensitivity to food-related odors in the human genome. Curr Biol 23, 1596–1600, doi:10.1016/j.cub.2013.07.031 (2013).

50 Zhuang, H. & Matsunami, H. Synergism of accessory factors in functional expression of mammalian odorant receptors. J Biol Chem 282, 15284–15293, doi:10.1074/jbc.M700386200 (2007).

51 Fukutani, Y. et al. The N-terminal region of RTP1S plays important roles in dimer formation and odorant receptor-trafficking. J Biol Chem 294, 14661–14673, doi:10.1074/jbc.RA118.007110 (2019).

52 Saito, H., Kubota, M., Roberts, R. W., Chi, Q. & Matsunami, H. RTP family members induce functional expression of mammalian odorant receptors. Cell 119, 679–691, doi:10.1016/j.cell.2004.11.021 (2004).

53 Yoshio, Y. & Nagata, E.

54 Tunyasuvunakool, K. et al. Highly accurate protein structure prediction for the human proteome. Nature 596, 590–596, doi:10.1038/s41586-021-03828-1 (2021).

55 Kim, S. et al. PubChem in 2021: new data content and improved web interfaces. Nucleic Acids Res 49, D1388–D1395, doi:10.1093/nar/gkaa971 (2021).

56 O’Boyle, N. M. et al. Open Babel: An open chemical toolbox. J Cheminform 3, 33, doi:10.1186/1758-2946-3-33 (2011).

57 Goodsell, D. S. & Olson, A. J. Automated docking of substrates to proteins by simulated annealing. Proteins 8, 195–202, doi:10.1002/prot.340080302 (1990).

58 Eberhardt, J., Santos-Martins, D., Tillack, A. F. & Forli, S. AutoDock Vina 1.2.0: New Docking Methods, Expanded Force Field, and Python Bindings. J Chem Inf Model 61, 3891–3898, doi:10.1021/acs.jcim.1c00203 (2021).

59 Trott, O. & Olson, A. J. AutoDock Vina: improving the speed and accuracy of docking with a new scoring function, efficient optimization, and multithreading. J Comput Chem 31, 455–461, doi:10.1002/jcc.21334 (2010).

60 Humphrey, W., Dalke, A. & Schulten, K. VMD: visual molecular dynamics. J Mol Graph 14, 33–38, 27-38, doi:10.1016/0263-7855(96)00018-5 (1996).

61 Pettersen, E. F. et al. UCSF Chimera--a visualization system for exploratory research and analysis. J Comput Chem 25, 1605–1612, doi:10.1002/jcc.20084 (2004).

